# Jacalin domain-containing protein *Os*SalT interacts with *Os*DREB2A and *Os*NAC1 to impart drought stress tolerance *in planta*

**DOI:** 10.1101/2020.04.19.049296

**Authors:** Salman Sahid, Chandan Roy, Dibyendu Shee, Riddhi Datta, Soumitra Paul

**Author notes:** Equal Contribution (Salman Sahid, Chandan Roy).

## Abstract

With the changing climatic conditions, drought has become one of the most threatening abiotic stress factors that adversely affect rice cultivation and productivity. Although the involvement of the jacalin domain-containing protein, *Os*SalT, has been reported in drought and salinity tolerance, its functional mechanism still remains largely unexplored. In this study, expression of the *OsSalT* gene was found to be positively correlated with the drought tolerance potential with its higher transcript abundance in the tolerant *indica* rice cultivar, Vandana and lower abundance in the susceptible cultivar, MTU1010. Moreover, the ectopic expression of *Os*SalT in tobacco imparted drought stress tolerance in the transgenic lines. The transgenic lines exhibited significantly improved growth and higher osmolyte accumulation over the wild-type (WT) plants together with the induction in the *Os*SalT expression under drought stress. Fascinatingly, the yeast two-hybrid and bimolecular fluorescence complementation (BiFC) analyses confirmed the interaction of *Os*SalT protein with two interesting transcription factors (TFs), *Os*NAC1 and OsDREB2A. *In silico* analysis further revealed that the *Os*SalT protein interacted with the regulatory domain of *Os*DREB2A and the C-terminal domain of *Os*NAC1 leading to their activation and induction of their downstream drought-responsive genes. Together, this study unravels a novel model for *Os*SalT-mediated regulation of drought tolerance in plants.

## 1. Introduction

Drought is among the most important abiotic stress factors that affect crop productivity resulting in significant yield losses (Hu and Xiang 2014). The changing climatic conditions like irregularity and insufficiency of rainfall adversely affect plant growth, development and productivity. Plants have evolved a highly sophisticated adaptive mechanism to combat drought stress. In-depth analysis of drought tolerance mechanism suggests an intricate crosstalk between different signalling cascades in plants. The drought stress response in plants is regulated by two distinct signalling pathways - the abscisic acid (ABA)-dependent and ABA-independent pathways. Several important transcription factors (TFs) such as myelocytomatosis oncogene (MYC), myeloblastosis oncogene (MYB), basic leucine zipper (bZIP), NAM, ATAF, and CUC (NAC) and dehydration responsive element binding (DREB) function in both the pathways and are highly accumulated in response to drought stress. These TFs are known to modulate downstream osmotic stress-responsive genes to render stress tolerance (Nakashima et al. 2009; Atkinson and Urwin 2012).

Lectins are a family of carbohydrate-binding proteins that specifically recognize sugar moieties. Lectins are known to play diverse roles in plant defence, in addition to its involvement in various other physiological phenomena. For example, lectin proteins were found to be induced in response to heat shock in cell suspension cultures (Spadoro-Tank and Etzler 1988; Shakirova et al. 1996). Again, transgenic Thai rice cultivars ectopically expressing snowdrop lectin exhibited resistance to sap-sucking insects (Tinjuangjun et al. 2000). ASAL, a novel lectin from garlic, provided resistance against phloem-limited viruses in rice (Saha et al. 2006). In addition, a large number of pattern recognition receptors in plants contain lectin domains that are essential for pathogen recognition and defence (Kaku et al. 2006; Bellande et al. 2017).

Plant lectins usually constitute a heterogeneous group that has been classified into 7 subfamilies based on their structure and evolutionary relatedness (van Damme et al. 1998). However, a later study has recognized 12 families among which 4 families encompassed the newly identified lectin members (Jiang et al. 2010). Among them, the jacalin-related lectin (JRL) subfamily is characterized by the presence of a conserved disaccharide-binding jacalin domain that was first identified in jackfruit (*Artocarpus integrifolia*) (de Azevedo Moreira and Ainouz 1981). The JRL subfamily comprises of at least 30 JRL proteins in rice distributed over its 9 out of 12 chromosomes. Among them 16 members contain only one jacalin domain and are called merojacalins. Thirteen members are chimeric jacalins containing at least one of the NB_ARC, Pkinase, dirigent or peptidase domains, in addition to the jacalin domain. The remaining one member has 3 repeats of the jacalin domain and is known as holojacalin (Yi-juan et al. 2018). The JRL proteins can again be categorized into two groups: the galactose-binding JRLs (gJRLs) with vacuolar localization and the mannose-binding JRLs (mJRLs) with nucleo-cytoplasmic localization (Van Damme et al. 2004; Lannoo and Van Damme 2014).

Members of the JRL subfamily have been demonstrated to play a crucial role in plant defence. Rice lectins containing mannose-binding jacalin domain have been demonstrated to play a role in salinity stress (Zhang et al. 2000). A map-based cloning strategy revealed that the *Jacalin-type Lectin Required for Potexvirus Resistance 1* (*JAX1*) gene confers resistance to *Plantago asiatica* mosaic virus infection in *Arabidopsis* (Yamaji et al. 2012). Chimeric JRLs have been reported to enhance tolerance against pathogen invasion, insect attack as well as abiotic stress in monocots (Song et al. 2014). Another jacalin like lectin protein, *Os*JAC1 imparted a high level of resistance against *Magnaporthe* infection in rice (Weidenbach et al. 2016).

An interesting JRL subfamily member, the *Os*SalT protein, is a 14.5 kDa mannose-binding lectin which comprises a single jacalin domain. It has been identified to accumulate in the root and sheath tissues of rice in response to salinity and drought stress (Claes et al. 1990; Filho et al. 2003). Transgenic rice plants overexpressing *OsSalT* gene have been shown to significantly improve salinity tolerance (He et al. 2016). In addition, a proteomic study has revealed that *Os*SalT was up-accumulated in rice roots under drought stress conditions (Paul et al. 2015). These pieces of evidence suggest a possible role of *Os*SalT protein in drought tolerance. However, the mechanism of how it imparts drought tolerance in plants has not been investigated so far.

In the present study, we have aimed to decipher the molecular mechanism of the drought-responsive role of *Os*SalT protein. Ectopic expression of *OsSalT* gene in tobacco significantly enhanced drought tolerance. We further identified that the *Os*SalT protein interacts with *Os*DREB2A and *Os*NAC1 TFs which induce the downstream drought-responsive gene expression thus imparting stress tolerance in plants.

## 2. Materials and Methods

### 2.1. Plant material, growth condition and stress assay

Seeds of *indica* rice cultivars, Vandana and MTU1010, were procured from the Chinsurah Rice Research Station, West Bengal, India. Seeds were germinated and grown in the net house under optimum conditions. The rice cultivars were normally irrigated for several weeks after transplanting. For stress assay, plants in the booting stage were exposed to drought by the withdrawal of watering for 7 days until the soil moisture content reached 45 % as standardized before (Paul et al. 2015).

Seeds of tobacco (*Nicotiana tabaccum* L. *cv. Xanthi*) were germinated and grown in Murashige and Skoog (MS) medium under 16 h light/8 h dark photoperiod as standardized before (Murashige and Skoog 1962; Ghanta et al. 2013). For stress assay, 60 days old soil-grown plants were subjected to drought stress by complete withdrawal of watering for 5 days until soil moisture content reached 45 %.

### 2.2. Morphological analysis

Different agronomic parameters such as plant height, percentage of brown leaves, percentage of rolled leaves, shoot and root dry weight were recorded for the two rice cultivars under control and drought conditions. For tobacco, plant height, percentage of yellow leaves, percentage of rolled leaves, shoot and root dry weight were recorded for wild-type (WT) and transgenic (OX) lines.

### 2.3. Biochemical analysis

For estimation of proline content, 0.2 g leaf tissue was collected from each sample and extracted with ethanol. The supernatant was mixed with 1% ninhydrin solution (in 60% acetic acid) and 20% ethanol and incubated at 95 °C for 30 min. The optical density was measured in spectrophotometer (Hitachi) at 595 nm (Woodrow et al. 2016). The leaf samples were homogenized in 0.05 % toluene and glycine betaine content was estimated following Grieve and Grattan (1983). Soluble sugar content was estimated according to Dubois (1956). Briefly, samples were homogenized in double distilled water, boiled at 100 °C for 30 min and centrifuged at 5000 rpm for 5 min. The supernatant was mixed with 9% phenol and 98% sulfuric acid, incubated at room temperature for 30 min and optical density was recorded at 485 nm. Total chlorophyll was extracted with 80 % acetone and quantified according to Lichtenthaler (1987).

### 2.4. RNA Isolation and q-RT PCR

Total RNA was isolated following Trizol method. Complementary DNA was synthesized using iScript cDNA synthesis kit (Bio-rad) following the manufacturer’s instructions. The qPCR analysis was performed in CFX96 Real-time PCR Detection System (Bio-rad) by using iTaq Universal SYBR Green Supermix (Bio-rad) and gene-specific primers (Table S1). *Actin* was used as the reference gene.

### 2.5. Cloning, vector construction and raising of transgenic lines

The *OsSalT* gene was cloned into pGEMT-Easy vector (Promega) followed by sub-cloning into the *EcoRI* and *BamHI* RE sites of pEGAD vector under control of *CaMV35S* promoter. The resulting construct, *CaMV35S::OsSalT-GFP*, was transformed into tobacco using *Agrobacterium*-mediated (*Agrobacterium tumefaciens* GV3101) leaf disc method as standardized before (Ghanta et al. 2013). The putative transgenic plants were screened through genomic DNA PCR with *bar* gene-specific primer. The positive transgenic lines were maintained up to T_2_ generation which was used for further experiments. The ectopic expression of the transgene was confirmed by qRT-PCR analysis using gene-specific primers.

### 2.6. Yeast two hybrid assay

Yeast two hybrid assay was performed using Matchmaker Gold Yeast Two-Hybrid System (Clontech). The *OsSalT* gene was cloned in-frame into the *EcoRI* and *BamHI* RE sites of the yeast vector, pGBKT7 to obtain a *BD-OsSalT* construct as bait. To study the protein-protein interaction of *OsSalT*, a cDNA library, ligated to pGADT7-*Rec* as prey, was prepared from rice roots under drought stress using Make Your Own “Mate and Plate” Library System (Clontech). The constructs were co-transformed into Y2H Gold yeast strain and screened on DDO (SD/-Leu/-Trp) and QDO (SD/–Ade/–His/–Leu/–Trp) plates. This was followed by a QDO/X/A (SD/–Ade/–His/–Leu/–Trp/X-α-Galactosidase/Aureobasidin-A) selection to test interaction. To identify the interacting protein partner(s), selected colonies were analysed by PCR followed by sequencing according to the manufacturer’s instruction. The *AD-T-antigen* and *BD-p53* interaction was used as a positive control.

### 2.7. Bimolecular fluorescence complementation (BiFC) assay

The *OsSalT* gene was cloned into the *35S::pVYCE-cVenus* vector between *SpeI* and *BamHI* RE sites to obtain *35S::OsSalT-cVenus* construct. The interacting partners, *OsDREB2A* and *OsNAC1* were cloned into *35S::pVYNE-nVenus* vector between the same RE sites to obtain *35S::OsDREB2A-nVenus* and *35S::OsNAC1-nVenus* constructs (Waadt et al. 2008). These constructs were introduced into the onion epidermal cells in pairwise combinations according to Yang et al. (2014). The *35S::OsSalT-cVenus* construct with empty *35S::pVYNE-nVenus* vector was used as negative control. Samples were examined under laser scanning confocal microscope (Olympus FV1000-IX81) using excitation wavelength of 514 nm.

### 2.8. Fold prediction and homology modelling of OsSalT, OsDREB2A and OsNAC1 proteins

The *Os*SalT (Uniprot ID: Q0JMY8), *Os*DREB2A (Uniprot ID: Q0JQF7) and *Os*NAC1 (Uniprot ID: Q8H0I5) sequences were subjected to PSIPRED (Buchan and Jones 2019), PHYRE2 (Kelley et al. 2015) and ROBETTA (Song et al. 2013) for fold prediction and homology modelling. The 3D structures of the protein sequences were filtered or energy minimized by YASARA server (Krieger et al. 2009). Models were further validated by PROCHECK (Laskowski et al. 1996) using the Ramachandran plot. According to Ramachandran plot, the residues of the disordered region was further refined by using MODELLER based ModLoop (Fiser and Sali 2003) webserver (only the loop regions of a 3D model were refined). The final models were re-validated by PROCHECK.

### 2.9. Molecular docking analysis

The best-fitted model of *Os*DREB2A and *Os*NAC1 were used for molecular docking with *Os*SalT through protein-protein docking server PatchDock (Schneidman-Duhovny et al. 2005) and GRAMM-X (Tovchigrechko and Vakser 2006). Top 100 dock solutions were generated by the PatchDock web server. Solutions were further refined and rescored using the FireDock docking refinement server (Mashiach et al. 2008). The best dock solution was selected according to its lowest global energy. Approximate ΔG of binding of best selected docked solutions was calculated by using the PDBePISA server (Krissinel and Henrick 2007).

### 2.10. Sequence analysis and prediction of ubiquitination and SUMOylation sites

The protein sequence for *Os*DREB2A was analyzed manually for the presence of NRD domain on the basis of existing reports (Mizoi et al. 2019). The DNA-binding domain residues were retrieved from NCBI database. Ubiquitination sites were predicted using UbiSite prediction server (http://csb.cse.yzu.edu.tw/UbiSite/prediction.php) with specificity level high. For SUMOylation consensus site prediction, GPS-SUMO web server (Ren et al. 2009; Zhao et al. 2014) was used with specificity level set as high.

### 2.11. Statistical Analysis

All experiments were performed in three independent biological replicates and data were represented as mean ± standard error of mean (SEM). Differences in morphological parameters, metabolite contents and transcript abundances among genotypes and treatments were assessed by two-way ANOVA followed by Sidak’s multiple comparison tests using the GraphPad Prism version 8.0.0 (GraphPad Software, San Diego, California USA, www.graphpad.com). The statistical significance at P ≤ 0.05 to identify the difference between the two sets of data was considered.

## 3. Results

### 3.1. Expression of OsSalT gene positively co-related with the drought tolerance potential of two rice cultivars

Two *indica* rice cultivars, Vandana and MTU1010 were analyzed for their drought tolerance potential. Several morphological parameters such as plant height, percentage of rolled and brown leaves, and root and shoot dry weights were estimated under control and drought conditions. No significant alteration in plant height and shoot dry weight was observed in response to drought stress in both the varieties. The root dry weight increased in Vandana under drought stress conditions while no alteration was observed for MTU1010 roots. The percentage of rolled and brown leaves was, however, significantly lower in the case of Vandana in comparison to MTU1010 under drought stress conditions (Fig. 1).

**Fig. 1.**
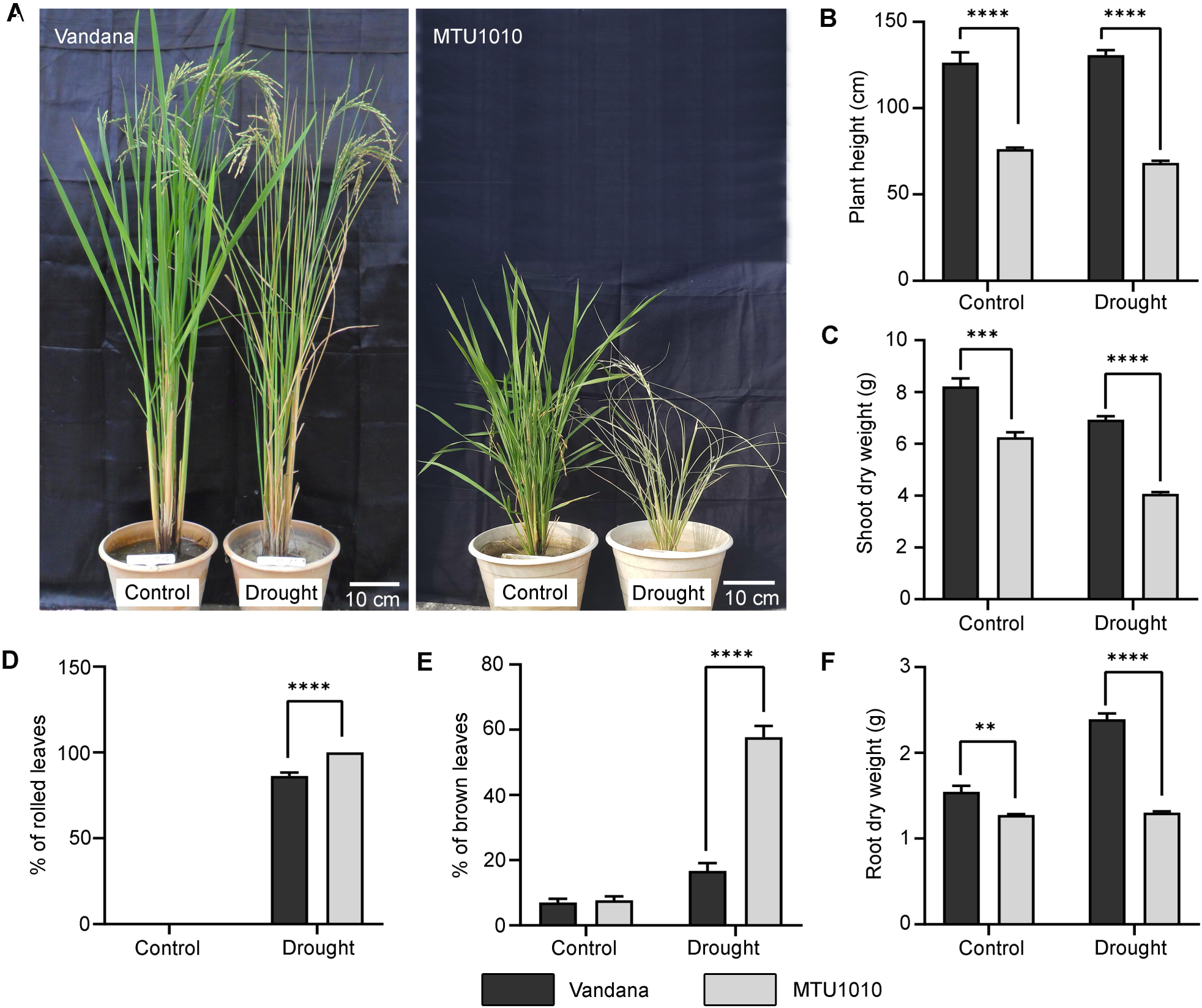
Drought stress analysis of two *indica* rice cultivars, Vandana and MTU1010. 100 days old plants were subjected to drought stress for 7 days and different morphological parameters were analyzed. (A) Morphology of Vandana and MTU1010 in response to drought stress, (B) Plant height, (C) Shoot dry weight, (D) Percentage of rolled leaves, (E) Percentage of brown leaves, and (F) Root dry weight. Results were represented as mean±SEM (n=3). Statistical difference between the cultivars under control and drought stress was denoted by asterisks at p<0.1 (**), p<0.01 (***) and p<0.001 (****).

To determine the effect of drought stress on the two cultivars, different osmotic stress-related metabolites such as proline, soluble sugars, glycine betaine, and chlorophyll contents were estimated. The total chlorophyll content decreased in response to drought stress in both the cultivars and the content was considerably lower in MTU1010 in comparison to Vandana. The proline and glycine betaine contents increased in response to drought stress in both the cultivars in a similar pattern. In case of soluble sugars, however, the content was more steeply increased in case of Vandana (97.29 mg/g FW to 158.8 mg/g FW) than in MTU1010 (55.7 mg/g FW to 92.49 mg/g FW) in response to drought stress (Fig. 2A-D). Together, these morphological and biochemical analyses indicate that Vandana is significantly more tolerant to drought stress in comparison to MTU1010.

**Fig. 2.**
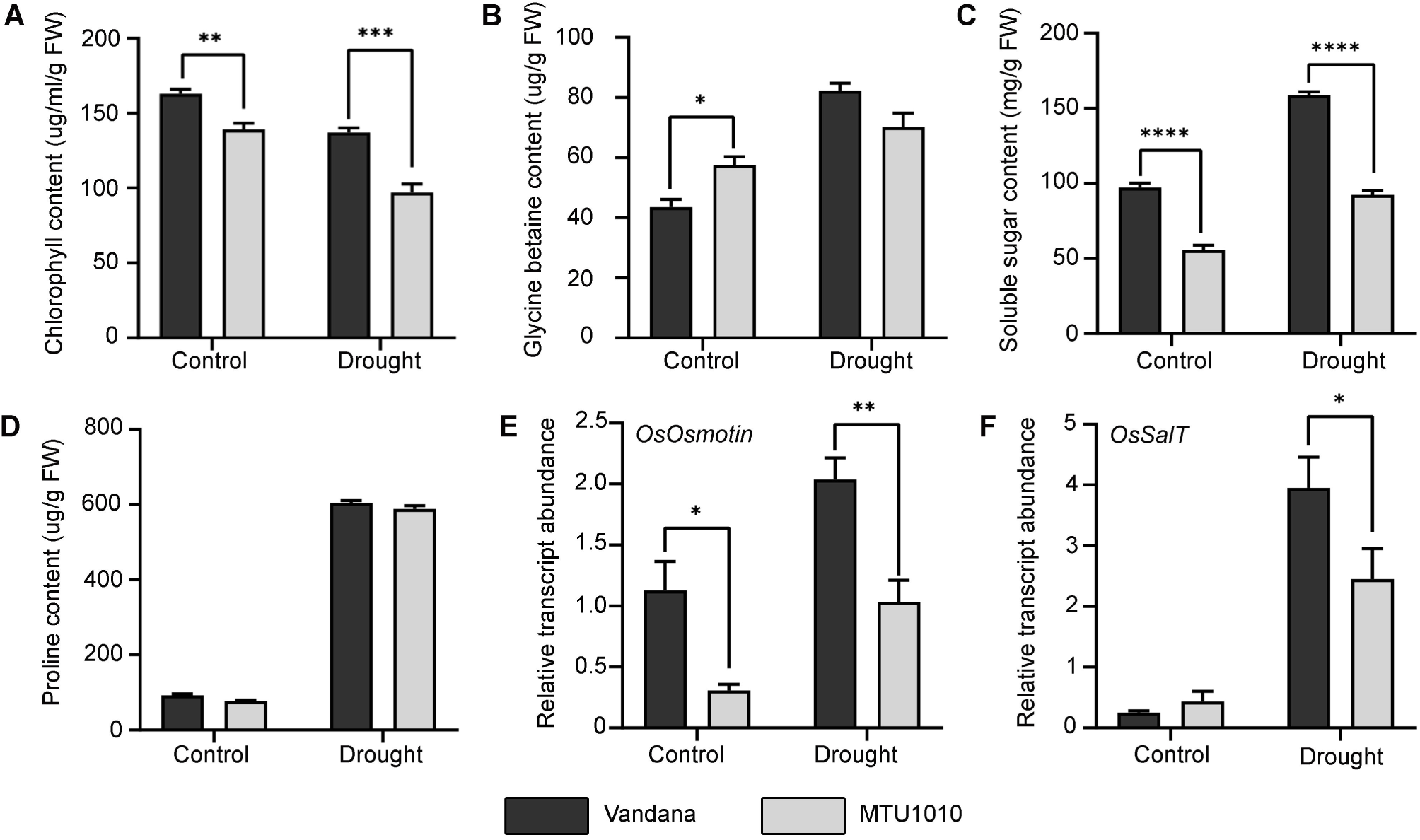
Biochemical and transcript analysis of Vandana and MTU1010 cultivars in response to drought stress. Samples were collected from control and stressed plants and several biochemical parameters were analyzed. (A) Chlorophyll content, (B) Glycine betaine content, (C) Soluble sugar content, and (D) Proline content. Samples were also used for qRT-PCR analysis to study the relative transcript abundance for (E) *OsOsmotin* and (F) *OsSalT* genes. Results were represented as mean±SEM (n=3). Statistical difference between the cultivars under control and drought stress was denoted by asterisks at p<0.05 (*), p<0.01 (**), p<0.001 (***) and p<0.0001 (****).

Furthermore, the expression of *OsSalT* along with *OsOsmotin*, a marker gene for drought stress, was analyzed in both Vandana and MTU1010 under drought stress. It was observed that the relative transcript abundance of *OsSalT* gene was increased by 15.61 fold in Vandana and by 5.61 fold in MTU1010 in response to drought stress. In the case of the *OsOsmotin* gene, the relative transcript abundance was significantly higher in Vandana in comparison to MTU1010 under both control and drought conditions (Fig. 2E-F). These observations suggested a positive correlation of *OsSalT* expression with the drought tolerance potential in rice.

### 3.2. Ectopic expression of OsSalT gene improved the drought tolerance in tobacco

To further validate the involvement of the *OsSalT* gene in imparting drought tolerance, the gene was ectopically expressed in tobacco. Among the 17 putative transformants generated, 11 lines were found to be positive (Fig. S1). Three best transgenic lines (OX1, OX2, and OX3) with the highest expression of *OsSalT* were considered for further experiments. Morphological analysis revealed no variations in these OX lines as compared to the WT and the vector control (VC) plants.

Stress assay was performed to assess the drought tolerance potential of the OX lines. It was observed that all the three OX lines were significantly more tolerant to drought stress in comparison to the WT and VC plants. The morphological analysis demonstrated that there was no alteration in shoot dry weight in the OX, VC and WT plants under control condition. However, the shoot dry weight was considerably higher in the case of the OX lines in comparison to the VC and WT plants under drought stress. The percentage of wilted leaves and yellow leaves were much higher in WT and VC plants than the OX lines (Fig. 3A-D).

**Fig. 3.**
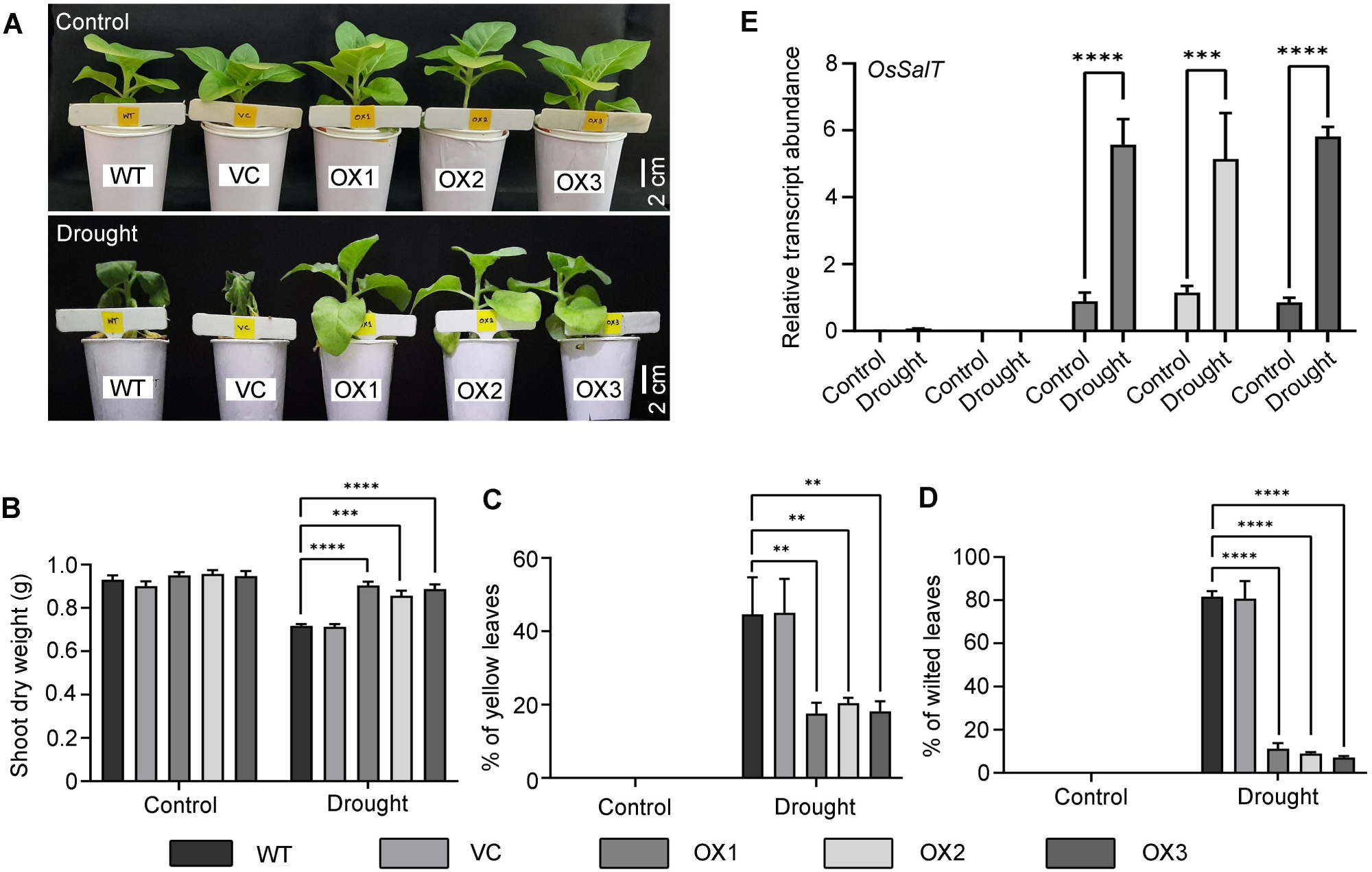
Response of transgenic tobacco lines ectopically expressing *OsSalT* gene under drought stress condition. The WT, VC and three independent transgenic lines (OX1, OX2 and OX3) were subjected to drought stress for 5 days and analyzed. (A) Morphology, (B) Shoot dry weight, (C) Percentage of yellow leaves and (D) Percentage of wilted leaves. The relative transcript abundance of OsSalT gene (E) was also analyzed. Results were represented as mean±SEM (n=3). Statistical difference between the lines under control and drought stress was denoted by asterisks at p<0.01 (**), p<0.001 (***) and p<0.0001 (****).

The accumulation of several osmotic stress-related metabolite contents under drought stress was also investigated. It was observed that the drop in the total chlorophyll content under drought stress was much steeper in case of the WT and VC plants in comparison to the OX lines. Accumulation of other metabolites like proline, glycine betaine and soluble sugars were also induced under drought stress condition. However, the induction was significantly higher in the OX lines than the WT and VC plants in response to drought stress (Fig. S2).

Besides, the expression of *OsSalT* in the WT, VC and OX lines were analyzed. As expected no expression was detected in the WT and VC plants. In the case of the OX1, OX2 and OX3 lines, the *OsSalT* expression was induced by 6.26, 4.47 and 6.76 folds respectively in response to drought stress (Fig. 3E). Together, these observations confirmed that the ectopic expression of *OsSalT* gene significantly improved drought stress tolerance in tobacco.

### 3.3. OsSalT protein interacts with OsDREB2A and OsNAC1 to impart drought tolerance

To explore the mechanism of how *OsSalT* imparts drought tolerance, the interacting protein partners were identified using yeast two-hybrid assay. A rice root library constructed under drought stress condition was used for this study. Out of 9 recombinant cDNA clones, 3 non-redundant putative interacting proteins partners were identified. Interestingly, two proteins, *Os*DREB2A and *Os*NAC1, were identified as positive independent interacting partners of *Os*SalT protein (Fig. 4A). The co-expression of *BD-OsSalT* with *AD-OsDREB2A* and *AD-OsNAC1* exhibited strong activation of X-α-galactosidase and aureobasidin A in the QDO/X/A medium. The interaction of *BD-p53* with *AD-T-antigen* was used as a positive control.

**Fig. 4.**
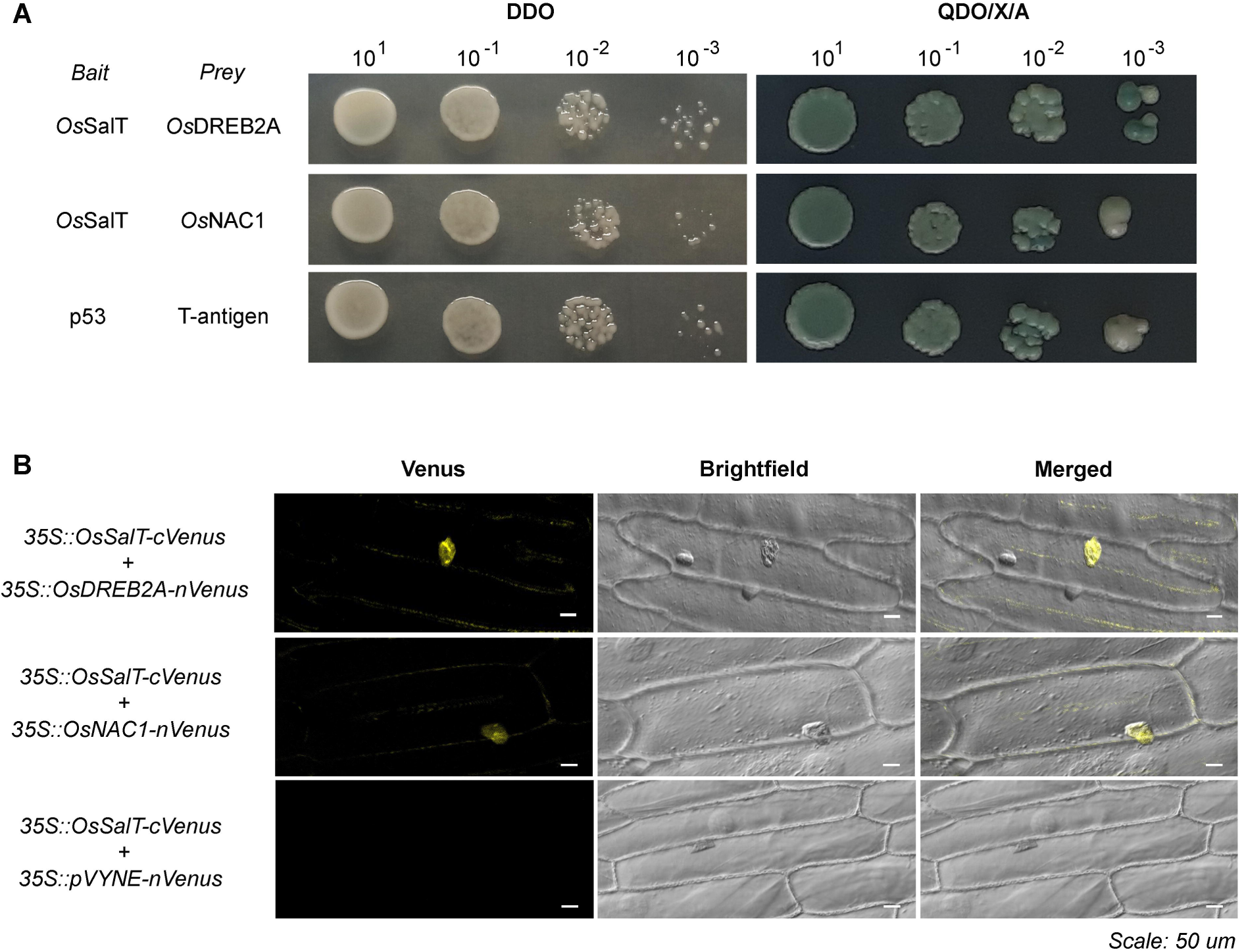
Identification of interacting protein partners of *Os*SalT protein under drought stress. (A) Yeast two-hybrid analysis identified the interaction of *Os*SalT with *Os*DREB2a and *Os*NAC1 proteins. The interaction of p53 protein with T-antigen was used as a positive control. (B) BiFC analysis confirmed the interaction of *Os*SalT protein with *Os*DREB2A in the nucleus and cytoplasm and with *Os*NAC1 in the nucleus only. Venus fluorescence, bright field, and merged images were represented for each set of constructs. Arrows indicate interaction in the nucleus.

Moreover, the interaction between *Os*SalT with *Os*DREB2A and *Os*NAC1 was confirmed by BiFC analysis (Fig. 4B). When the onion epidermal cells were co-transfected with the *35S::OsSalT-cVenus* and *35S::OsDREB2A-nVenus* constructs, strong yellow fluorescence was observed in the nucleus thus confirming a nuclear interaction. In the case of the *35S::OsSalT-cVenus* and *35S::OsNAC1-nVenus* constructs, yellow fluorescence was also detected in the nucleus indicating a nuclear interaction. However, no fluorescence was detected when the *35S::OsSalT-cVenus* construct was used with the empty *35S::pVYNE-nVenus* vector. These results confirmed the physical interaction of the *Os*SalT protein with *Os*DREB2A and *Os*NAC1 to regulate drought tolerance in plants.

### 3.4. In silico analysis confirmed the physical interaction of OsSalT protein with OsDREB2A and OsNAC1

The 3D structures for *Os*SalT, *Os*DREB2A and *Os*NAC1 proteins were generated by homology modelling followed by energy minimization. The quality of the models was assessed using PROCHECK server. Ramachandan plot of the complete structures revealed that 91.2 %, 90.3% and 87.6 % residues were present in the most favoured regions in case of the *Os*SalT, *Os*DREB2A and *Os*NAC1 proteins respectively. None of the residues were present in the disallowed region (Fig. S3-S8). Protein-protein interaction for *Os*SalT-*Os*DREB2A and *Os*SalT-*Os*NAC1 was studied *in silico* through molecular docking using PatchDock and GRAMM-X servers. The docking results confirmed physical interaction of the *Os*SalT protein with *Os*DREB2A and *Os*NAC1 proteins (Fig. 5-6). Sequence analysis along with the molecular docking data indicated that *Os*NAC1 protein possesses its DNA-binding domain at the N-terminal end while it interacts with the *Os*SalT protein at its C-terminal domain (Fig. S9). The docking analysis further revealed that the *Os*SalT protein binds to a regulatory region of the *Os*DREB2A protein without blocking its DNA-binding domain (residues 75-132) (Fig. 5, S10). This interaction presumably activates the *Os*DREB2A protein under drought stress thus enabling it to induce the transcription of the DREB-responsive genes.

**Fig. 5.**
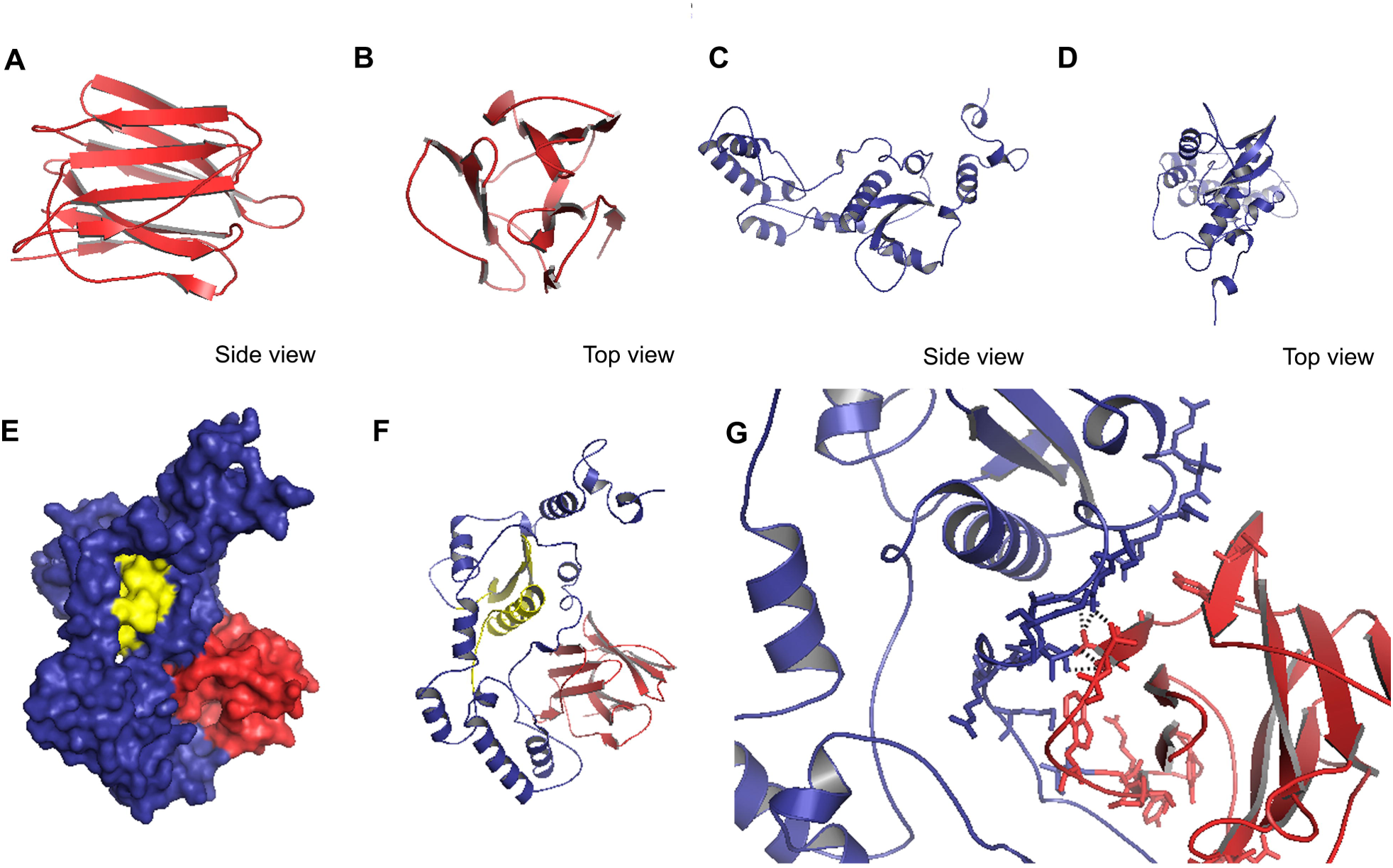
*In silico* analysis for *Os*SalT-*Os*DREB2A interaction. Protein structures for *Os*SalT and *Os*DREB2A were generated through homology modelling. The top views (A,C) and the side views (B,D) of *Os*SalT and *Os*DREB2A structures have been shown. The structures were used for molecular docking analysis. (E) Surface view, (F) Cartoon view, (G) Interphase residues. Black dashed lines indicate polar contacts among the interphase residues. Yellow residues indicate the DNA-binding domain of *Os*DREB2A.

**Fig. 6.**
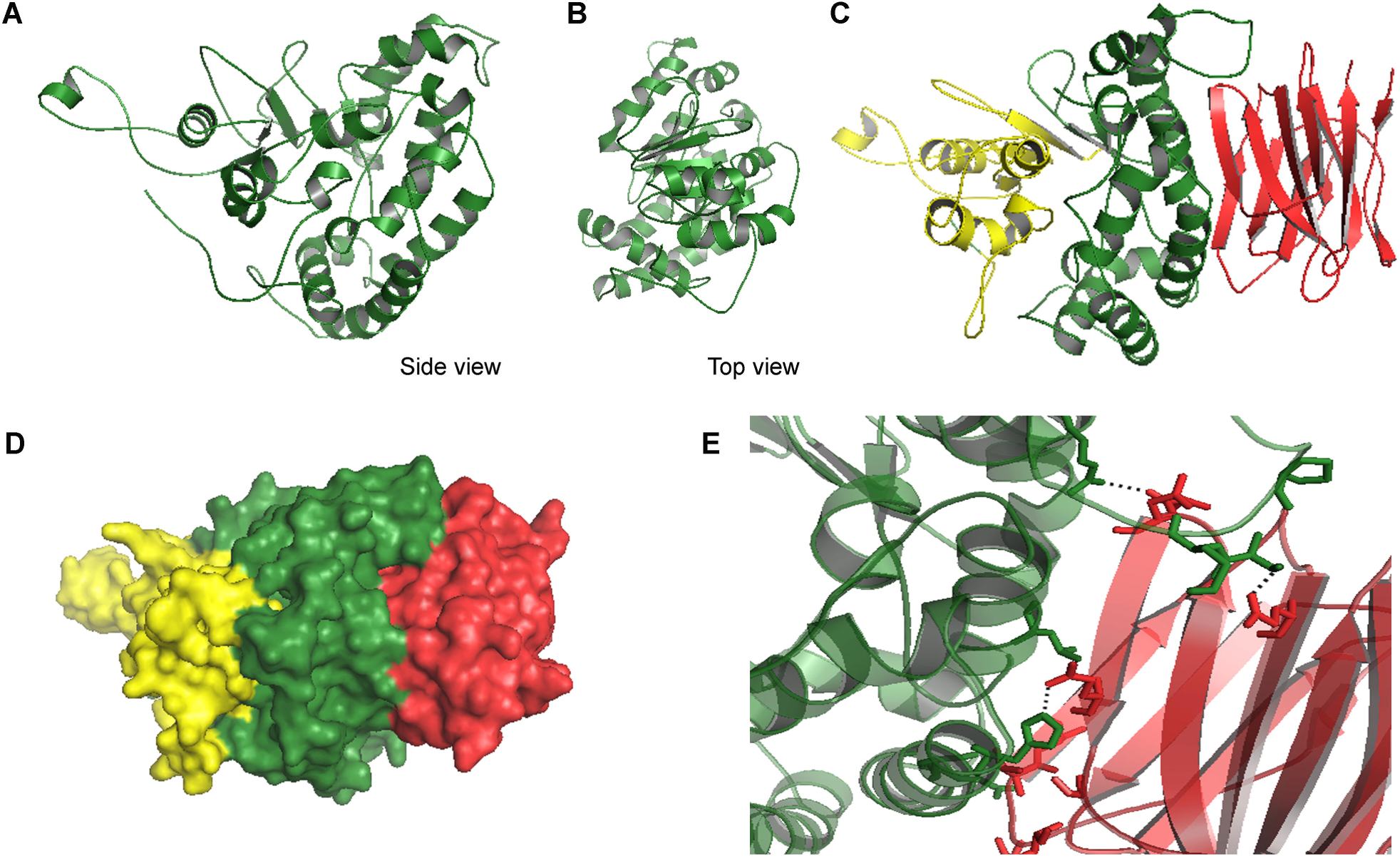
*In silico* analysis for *Os*SalT-*Os*NAC1 interaction. Protein structure for *Os*NAC1 was generated through homology modelling. The top view (A) and the side view (B) of *Os*NAC1structure have been shown. The structure was used for molecular docking analysis with *Os*SalT protein. (C) Cartoon view, (D) Surface view, (E) Interphase residues. Black dashed lines indicate polar contacts among the interphase residues. Yellow residues indicate the DNA-binding domain of *Os*NAC1.

It has been reported that the NRD domain of most DREB proteins contain an acidic sequence containing a stretch of at least five serine/threonine residue cluster (Mizoi et al. 2019). Analysis of the protein sequence revealed that *Os*DREB2A protein, unlike *Os*DREB2B and most DREB proteins in other plants, lacks a well-defined NRD regulatory region indicating that it may possess unique regulatory sites. Next, its structure was analyzed for probable ubiquitination and SUMOylation sites. It was observed that the protein possess three probable ubiquitination sites at K184, K185 and K199 (Fig. S10). In addition, a SUMOylation consensus site at K199 had been predicted which indicated a probable regulation via ubiquitination-SUMOylation switch (Fig. S10). It was further observed that this K199 does not overlap with both the DNA-binding as well as the *Os*SalT-interacting sites. These observations suggested a unique regulatory mechanism for *Os*DREB2A protein in rice.

### 3.5. Expression of DREB2A and NAC1 responsive genes were enhanced in transgenic tobacco plants (OX) under drought stress

To further confirm the *Os*SalT-mediated activation of DREB2A and NAC1 proteins, expression of several stress-responsive genes were analyzed under drought condition in the OX plants. It is known that the transcriptions of the *Dehydrin* and *Late Embryogenesis Abundant* (LEA) genes are induced by DREB2A while the *Phosphatidylinositol-3-kinase* (*PI3K*) and *Sucrose phosphate synthase* (*SPS*) are NAC1-responsive genes (Lata and Prasad 2011; Saad et al. 2013). Therefore, these genes were selected in this analysis. In addition, *Responsive to dehydration 22* (*rd22*) and *Alcohol dehydrogenase 1* (*ADH1*) genes, which function in DREB2A and NAC1 independent manner, were selected as negative control (Abe et al. 2003). Interestingly, it was observed that the expression of *Dehydrin, LEA, PI3K* and *SPS* genes were significantly enhanced in the OX plants under drought stress as compared to the WT plants (Fig. 7). On the contrary, expression of both *rd22* as well as *ADH1* genes was not significantly altered. This confirmed that *Os*SalT activates both DREB2A and NAC1 proteins that in turn results in the induction of the downstream drought-responsive genes ultimately leading to stress tolerance.

**Fig. 7.**
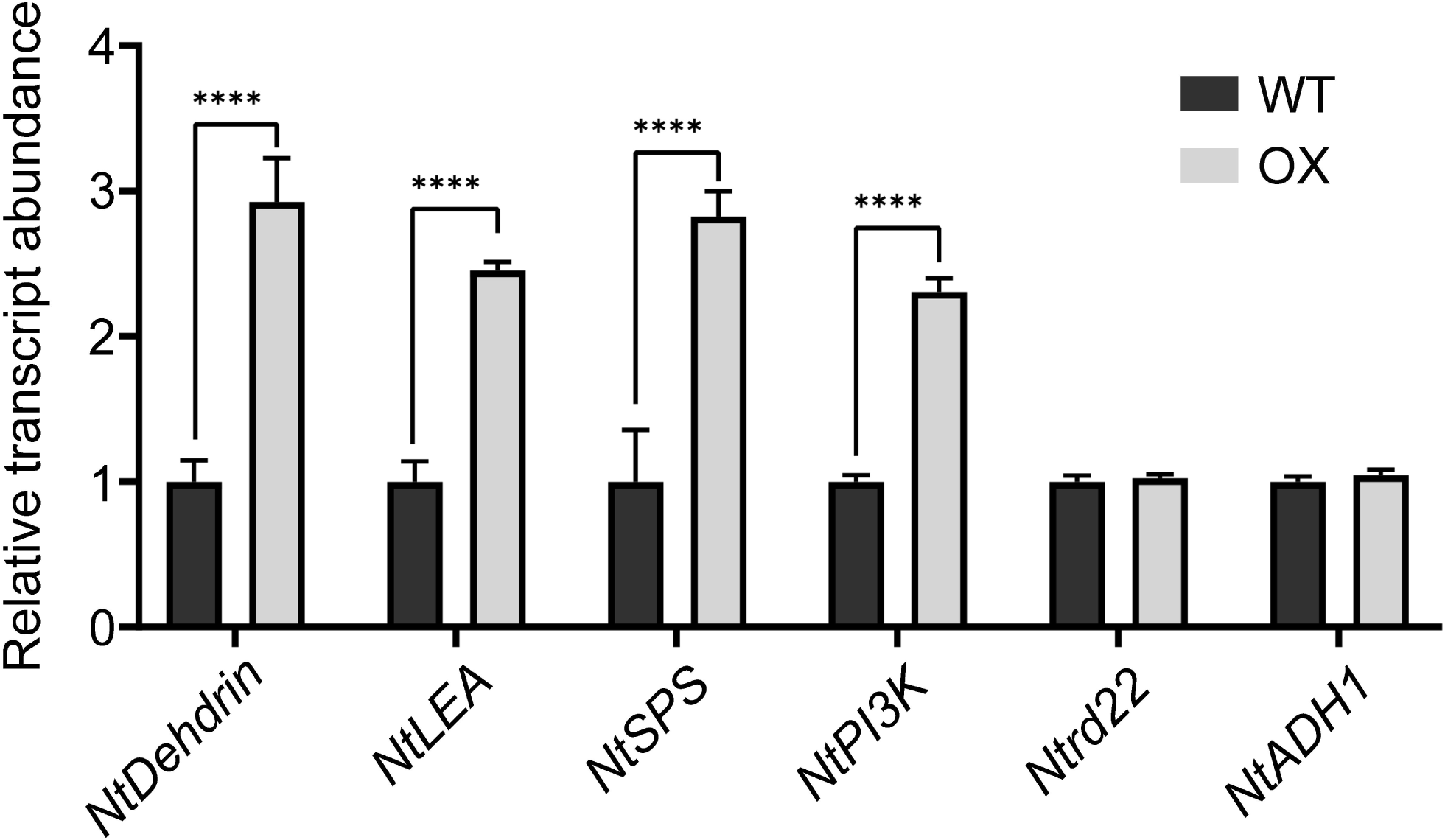
Expression of *DREB2A* and *NAC1* responsive genes under drought stress in the WT and transgenic tobacco (OX) plants. The relative transcript abundance of *NtDehydrin, NtLEA, NtSPS, NtPI3K, Ntrd22 and NtADH1* genes were analyzed under drought stress. Results were represented as mean ± SEM (n=3). Statistical difference between the lines under control and drought stress was denoted by asterisks at p<0.0001 (****).

## 4. Discussion

In plants, lectin proteins are widely known to be involved in various biotic and abiotic stress responses. Jacalin is a mannose-binding lectin protein that serves an important role in regulating blast disease and osmotic stress tolerance in rice (Zhang et al. 2000; Song et al. 2014). In this study, we have dissected the role of *Os*SalT, a jacalin domain containing lectin protein, in drought stress tolerance in plants. Earlier, it had been identified that the expression of *OsSalT* was augmented in rice in response to salt stress. The *OsSalT* overexpressing rice plants also displayed enhanced osmotic stress tolerance (Filho et al. 2003; He et al. 2016). However, how this *Os*SalT protein provides osmotic stress tolerance in plants remains elusive so far. Here, we have explored the molecular mechanism of *Os*SalT protein in regulating drought stress tolerance *in planta*.

To identify the drought stress-responsive role of *Os*SalT protein, its expression was analyzed in response to drought stress in a drought tolerant (Vandana) and a drought susceptible (MTU1010) *indica* rice cultivar. The expression of the *OsSalT* gene was found to be significantly higher in the drought-tolerant cultivar in comparison to the susceptible one under drought stress. This observation suggested a direct correlation of the *OsSalT* expression with the degree of drought tolerance in rice. Furthermore, the ectopic expression of *OsSalT* gene provided enhanced drought tolerance in tobacco. The OX plants displayed an improved wilting percentage and less chlorophyll degradation. The higher accumulation of several osmotically active compounds has long been associated with osmotic stress tolerance potential in plants (Chen and Murata 2002). In the present study, the OX lines exhibited an elevated accumulation of proline, glycine betaine and soluble sugar contents under drought stress which indicated their better adaptability over the WT plants.

With this confirmation, we aimed to decipher the molecular mechanism of how the *Os*SalT protein imparts drought tolerance in plants. It was identified that the *Os*SalT protein interacts with two interesting protein partners, *Os*DREB2A and *Os*NAC1 in rice roots under drought stress conditions. It has been widely demonstrated that both of these protein partners function as important TFs and can regulate the expression of different drought and thermotolerance-associated genes (Liu et al. 1998; Dubouzet et al. 2003; Saad et al. 2013). It was further observed that its interaction with both *Os*DREB2A and *Os*NAC1 occurs in the nucleus.

Emerging evidences suggested that NAC family comprises of a large group of plant-specific TFs named upon 3 protein members, NAM, APAF1-2, and CUC2. They play a pivotal role in several abiotic stress responses in plants (Aida et al. 1997). An array of NAC family members have been identified from different plant species including rice. In rice, 151 NAC family genes were reported while most of them were characterized as stress-responsive NAC genes (sNAC) (Nuruzzaman et al. 2010). Recently, the drought tolerance activity of *OsNAC1* has been confirmed in bread wheat (Saad et al. 2013). In the present study, it was identified that the jacalin domain-containing protein, *Os*SalT, can bind to the *Os*NAC1 protein in rice in response to drought stress. The *Os*NAC1 protein was found to possesses an N-terminal DNA binding domain which binds to several drought- or salinity-responsive genes (Aida et al. 1997). On the other hand, the *Os*SalT protein was found to interact with its C-terminal end to modulate its function. This observation can be corroborated with the previous report that all NAC proteins contain a highly conserved N-terminal DNA-binding domain and a variable C-terminal regulatory region (Kikuchi et al. 2000; Ooka et al. 2003). This study delineates a probable mechanism of *Os*NAC1 regulation in response to drought stress in rice.

Previous investigations highlighted that several *DREB* subfamily members regulate osmotic stress tolerance in plants (Dubouzet et al. 2003; Lata and Prasad 2011). It has been demonstrated that these DREB proteins are specifically regulated via modulation of their protein stability. Although the induction of *Arabidopsis DREB2A* gene expression was found to be significant under drought stress, the activation of downstream signalling was completely dependent on the stability of the DREB2A protein. This protein stability has been reported to be maintained by phosphorylation and ubiquitination at a regulatory NRD domain (Shinozaki and Yamaguchi-Shinozaki 2000; Sakuma et al. 2002; Yamaguchi-Shinozaki and Shinozaki 2006; Mizoi et al. 2019). Recently, it has been shown that SUMOylation at the K163 amino acid residue close to the NRD domain inhibited the ubiquitination during stress and stabilised the protein thus imparting thermotolerance in *Arabidopsis* (Wang et al. 2020). However, the mechanism of its activation during stress still remains unanswered. In addition, most of the studies dissecting the DREB2A regulation have been restricted to *Arabidopsis* and no detailed activation mechanism for *Os*DREB2A protein during drought stress has been undertaken.

Interestingly, the *Os*DREB2A protein in rice, unlike the *Os*DREB2B, does not possess any conserved NRD domain and the activation mechanism has not been studied so far. This motivated us to explore the *Os*DREB2A structure through *in silico* analysis. It was observed that the protein still possesses three probable ubiquitination sites at K184, K185 and K199 among which K199 is also a SUMOylation consensus site. This observation suggested that in spite of lacking a conserved NRD domain, the *Os*DREB2A protein can still be regulated by an ubiquitination-SUMOylation switch. The balance between these two phenomena maintains the *Os*DREB2A protein accumulation in the cell to a basal level under control condition while quickly augmenting its level under drought stress.

In addition, *Os*SalT interacted with *Os*DREB2A protein at a regulatory domain, other than the DNA-binding domain and the K199 residue. This presumably activated the protein to regulate its downstream drought stress-responsive genes. *Os*DREB2A is a TF that can bind to the dehydrin responsive element (DRE) motif of different drought stress-responsive genes like *Dehydrin* and *LEA* to regulate drought stress tolerance (Lata and Prasad 2011). In this study, it was also observed that the DREB2A responsive genes, *Dehydrin* and *LEA* were significantly up-regulated in the tobacco OX lines suggesting that *Os*SalT presumably activates the DREB2A protein in tobacco which in turn activates the downstream drought-responsive genes. In fact, a proteomic study of *Os*DREB2A overexpressing rice plants identified up-accumulation of the *Os*SalT protein in roots under drought stress which corroborates with the present observation (Paul et al. 2015).

In summary, the present investigation has demonstrated the functional role of *Os*SalT protein in imparting drought stress tolerance via its interaction with *Os*DREB2A and *Os*NAC1 proteins (Fig. 8). It can be hypothesized that under drought stress, the ubiquitination of *Os*DREB2A is blocked due to SUMOylation of a specific lysine residue. This inhibits *Os*DREB2A degradation thus augmenting its accumulation in the cell. The *Os*SalT protein then interacts with the stabilized *Os*DREB2A. The binding of *Os*SalT protein probably leads to a conformational change in the *Os*DREB2A protein which facilitates its activation under drought stress. It can then bind to the DRE sequence of the drought stress responsive genes leading to their transcriptional activation. On the other hand, drought stress induces the expression of *Os*NAC1. The *Os*SalT protein then interacts with and activates *Os*NAC1 protein which in turn induces the expression of its downstream drought-responsive genes. Together, the present study has revealed the novel role of a jacalin domain containing *Os*SalT protein in regulating two crucial TFs thus enhancing the drought stress tolerance *in planta*.

**Fig. 8.**
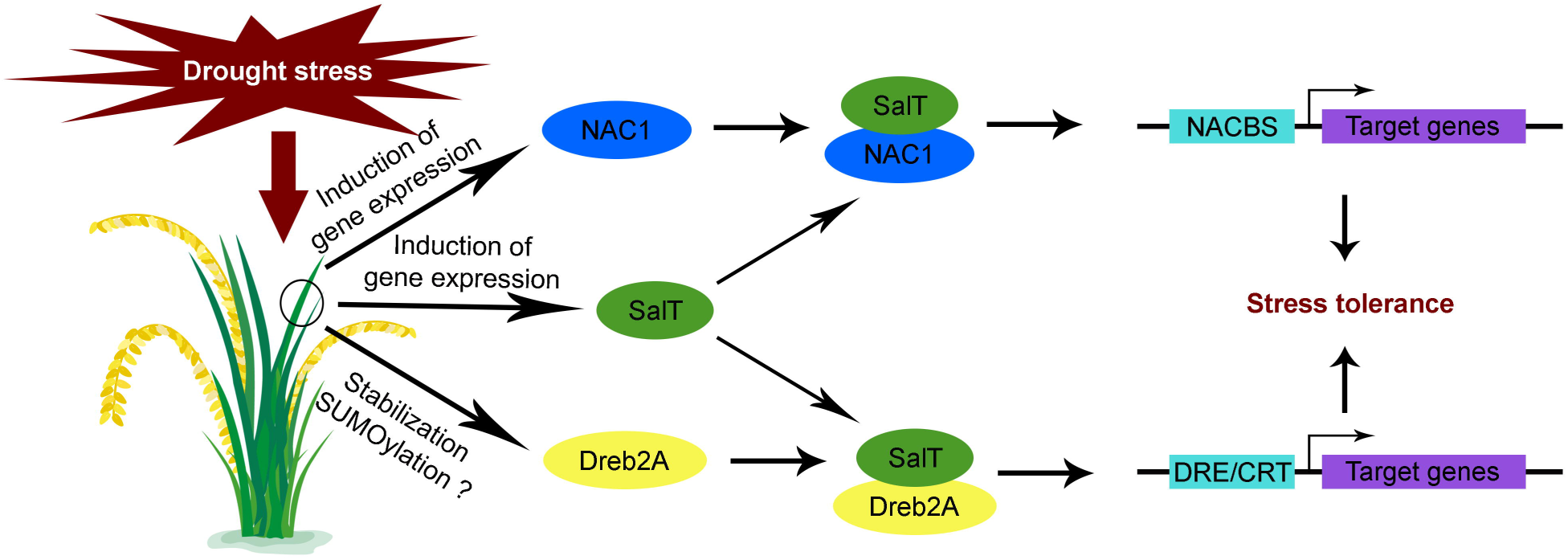
Model for *Os*SalT-mediated regulation of drought stress. Drought stress induces the expression of *Os*SalT in cells. In the mean tine, *Os*NAC1 expression is also induced and its accumulation increases. On the other hand, *Os*DREB2A protein is presumably stabilized by SUMOylation. The *Os*SalT now binds with the stabilized *Os*DREB2A and *Os*NAC1 proteins thus activating them. This results in the transcriptional activation of downstream drought-responsive genes leading to stress tolerance.

## Supporting information

ESM_1

ESM_2

## Author Contributions

RD and SP conceptualized and designed the research plan; SS and CR performed most of the experiments; DS performed the *in silico* analysis; RD and SP analyzed the data; SS drafted the manuscript; RD and SP supervised and complemented the writing.

## Acknowledgement

We thank the central instrumentation facilities of Department of Botany, University of Calcutta and Department of Botany, Dr. A. P. J. Abdul Kalam Government College as well as the confocal microscopic facility of DBT-IPLS, Department of Biochemistry, University of Calcutta. We thank Prof. Jörg Kudla (University of Munster, Germany) for providing the *pVYNE* and *pVYCE* vectors and Prof. Prabod Trivedi (CSIR-National Botanical Research Institute, India) for kindly sharing the *Agrobacterium tumefaciens* GV3101 strain.

## Supplementary data

**Supplementary Fig. S1. Screening of putative transformants by genomic DNA PCR for *bar* gene**. A prominent band at 412 bp indicated positive transgenic lines. L: 1 kb DNA ladder, N: negative control (WT), P: positive control (recombinant plasmid), 1-17: putative transformants.

**Supplementary Fig. S2: Biochemical analysis of transgenic tobacco lines ectopically expressing OsSalT gene under drought stress condition**. (A) Chlorophyll content, (B) Proline content, (C) Glycine betaine content, and (D) Soluble sugar content. Results were represented as mean±SEM (n=3). Statistical difference between the cultivars under control and drought stress was denoted by asterisks at p<0.0001 (****).

**Supplementary Fig. S3: Homology modelling of *Os*SalT protein**. (A) Fold prediction using PSIPRED, Surface view of the *Os*SalT protein in (B) side view and (C) top view.

**Supplementary Fig. S4: Ramachandran plot for *Os*SalT structure**.

**Supplementary Fig. S5: Homology modelling of *Os*DREB2A protein**. (A) Fold prediction using PSIPRED, Surface view of the *Os*DREB2A protein in (B) side view and (C) top view.

**Supplementary Fig. S6: Ramachandran plot for *Os*DREB2A structure**.

**Supplementary Fig. S7: Homology modelling of *Os*NAC1 protein**. (A) Fold prediction using PSIPRED, Surface view of the *Os*NAC1 protein in (B) side view and (C) top view.

**Supplementary Fig. S8: Ramachandran plot for *Os*NAC1 structure**.

**Supplementary Fig. S9: Sequence analysis for *Os*NAC1 protein showing the DNA-binding and OsSalT-interacting sites**.

**Supplementary Fig. S10: Sequence analysis for *Os*DREB2A protein**. (A) Protein sequence showing the DNA-binding, OsSalT-interacting, ubiquitination and SUMOylation consensus sites. (B) Ubiquitination site prediction. (C) SUMOylation site prediction.

**Supplementary Table S1: List of primers used**.

