## Supplementary material for "Jacalin domain-containing protein *Os*SalT interacts with *Os*DREB2A and *Os*NAC1 to impart drought stress tolerance *in planta*": ESM_1

*****Authors for correspondence:

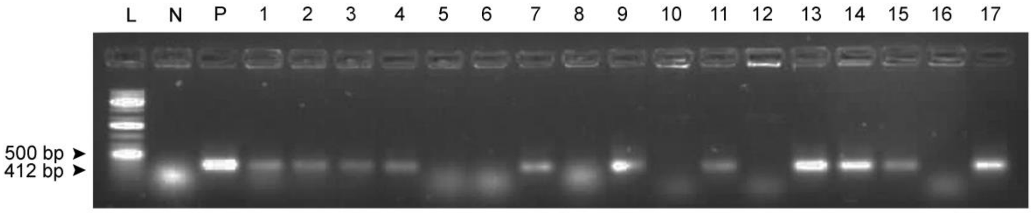


**Supplementary Fig. S1. Screening of putative transformants by genomic DNA PCR for *bar* gene.** A prominent band at 412 bp indicated positive transgenic lines. L: 1 kb DNA ladder, N: negative control (WT), P: positive control (recombinant plasmid), 1-17: putative transformants.

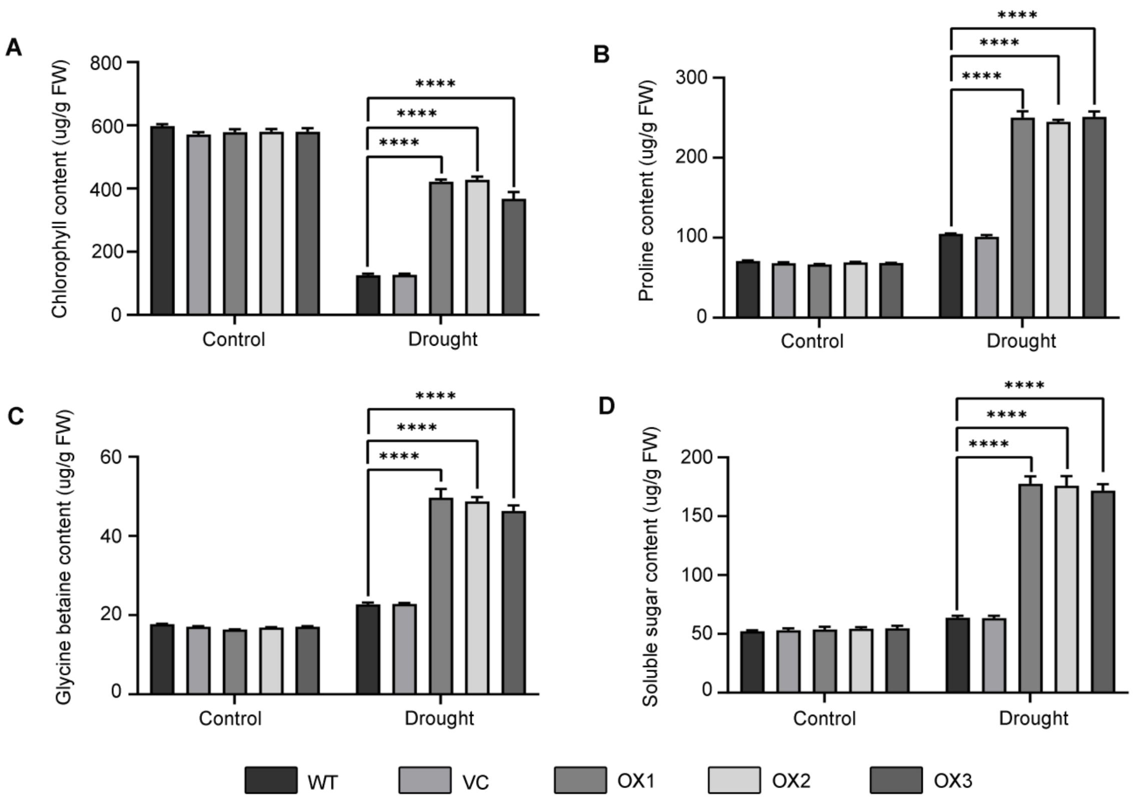


**Supplementary Fig. S2: Biochemical analysis of transgenic tobacco lines ectopically expressing OsSalT gene under drought stress condition.** (A) Chlorophyll content, (B) Proline content, (C) Glycine betaine content, and (D) Soluble sugar content. Results were represented as mean±SEM (n=3). Statistical difference between the cultivars under control and drought stress was denoted by asterisks at $p<0.0001$ (****).

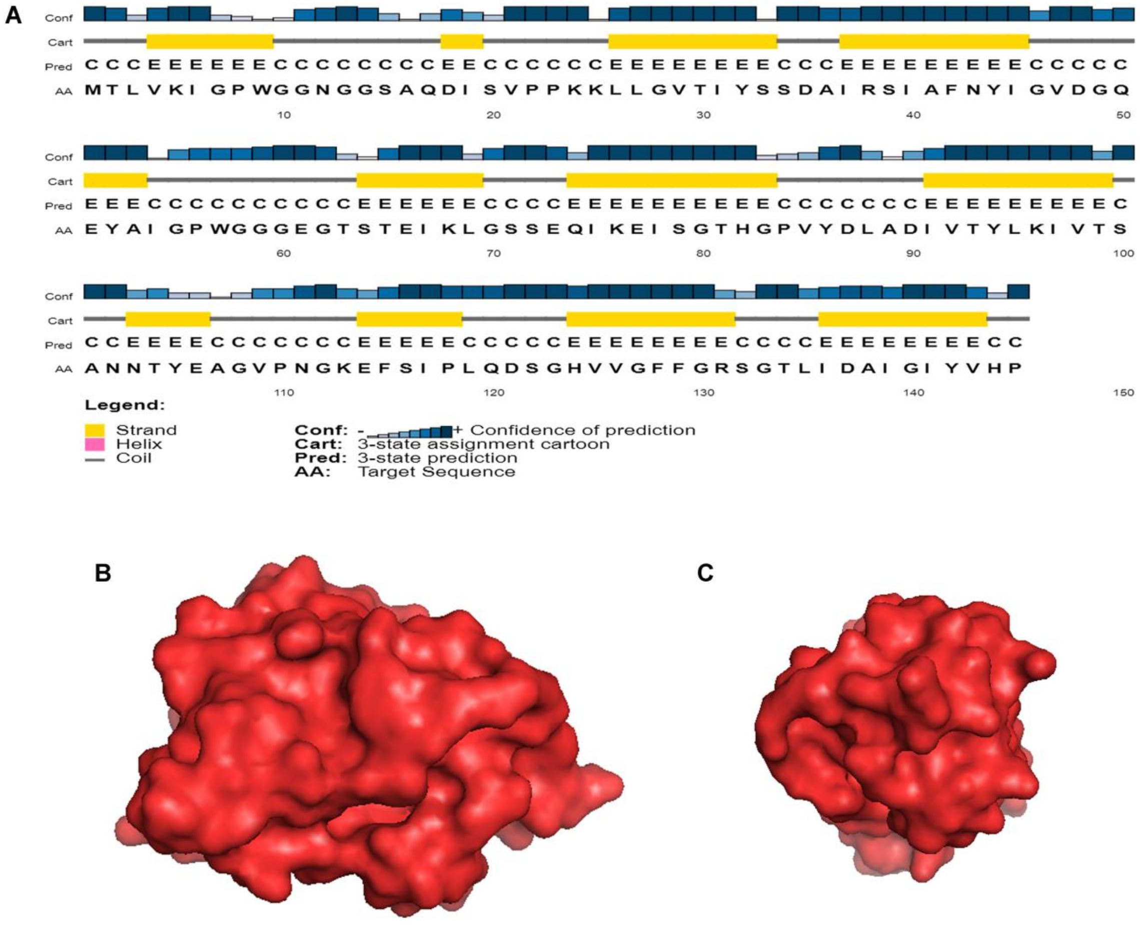


**Supplementary Fig. S3: Homology modelling of *Os*SalT protein.** (A) Fold prediction using PSIPRED, Surface view of the *Os*SalT protein in (B) side view and (C) top view.

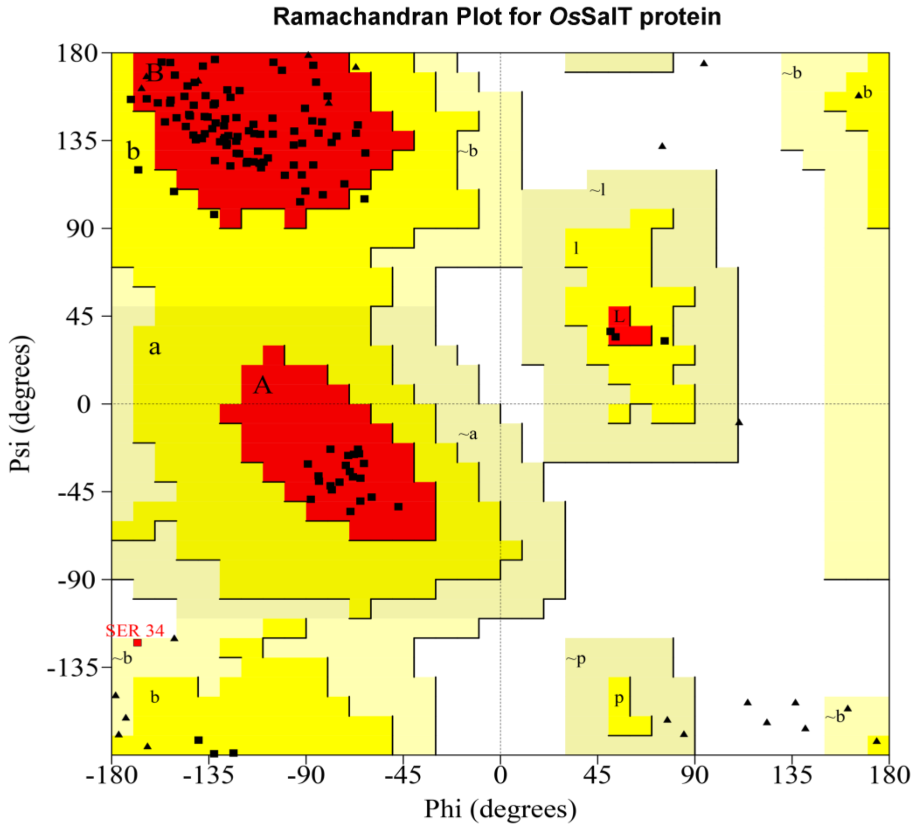


Plot statistics

| | | |
|---|---|---|
| Residues in most favoured regions [A,B,L] | 103 | 91.2% |
| Residues in additional allowed regions [a,b,l,p] | 9 | 8.0% |
| Residues in generously allowed regions [~a,~b,~l,~p] | 1 | 0.9% |
| Residues in disallowed regions | 0 | 0.0% |
|  | ---- | ------ |
| Number of non-glycine and non-proline residues | 113 | 100.0% |
| Number of end-residues (excl. Gly and Pro) | 1 | |
| Number of glycine residues (shown as triangles) | 23 | |
| Number of proline residues | 8 | |
|  | ---- |  |
| Total number of residues | 145 | |

**Supplementary Fig. S4: Ramachandran plot for *Os*SalT structure.**

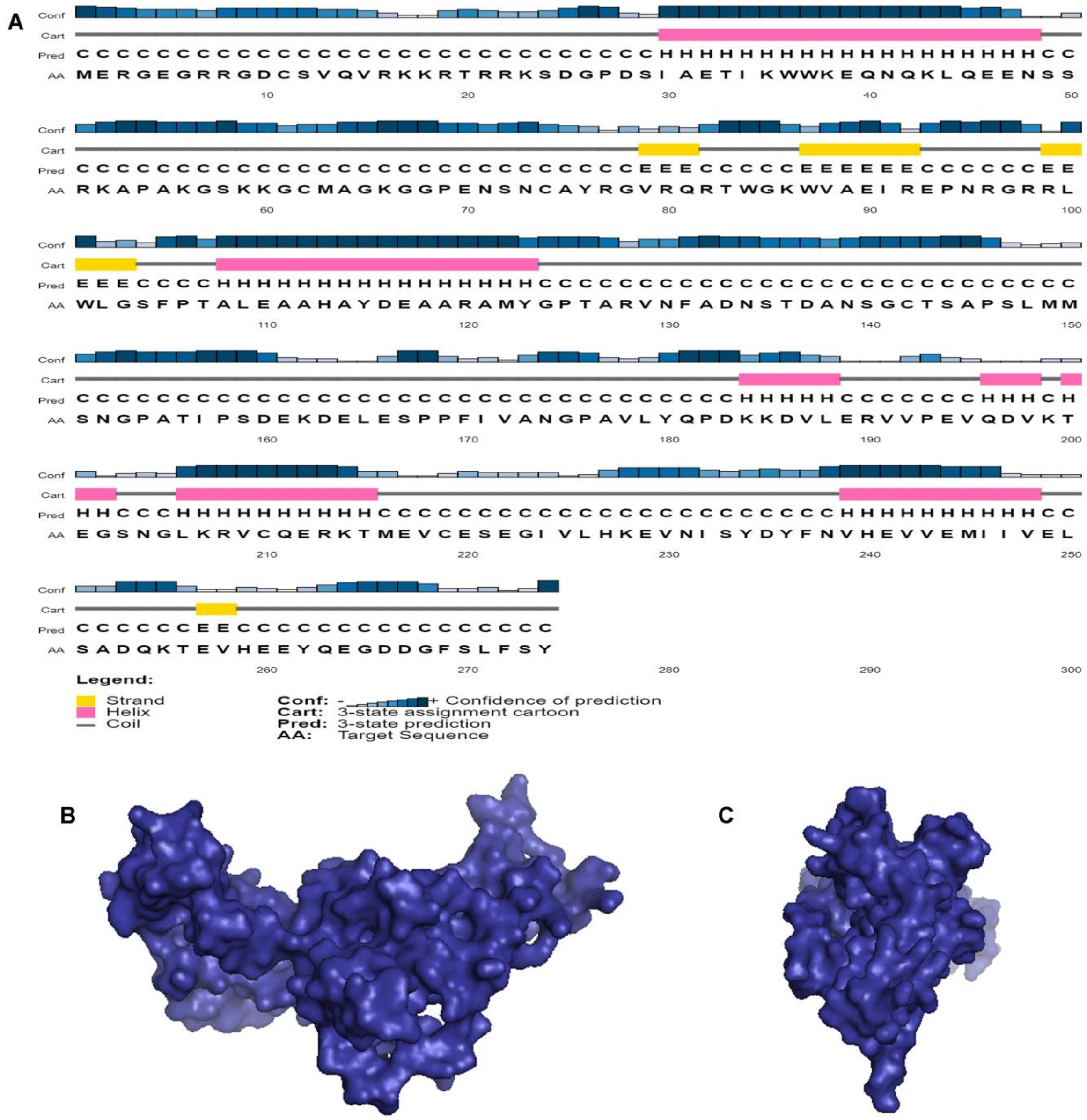


**Supplementary Fig. S5: Homology modelling of *Os*DREB2A protein.** (A) Fold prediction using PSIPRED, Surface view of the *Os*DREB2A protein in (B) side view and (C) top view.

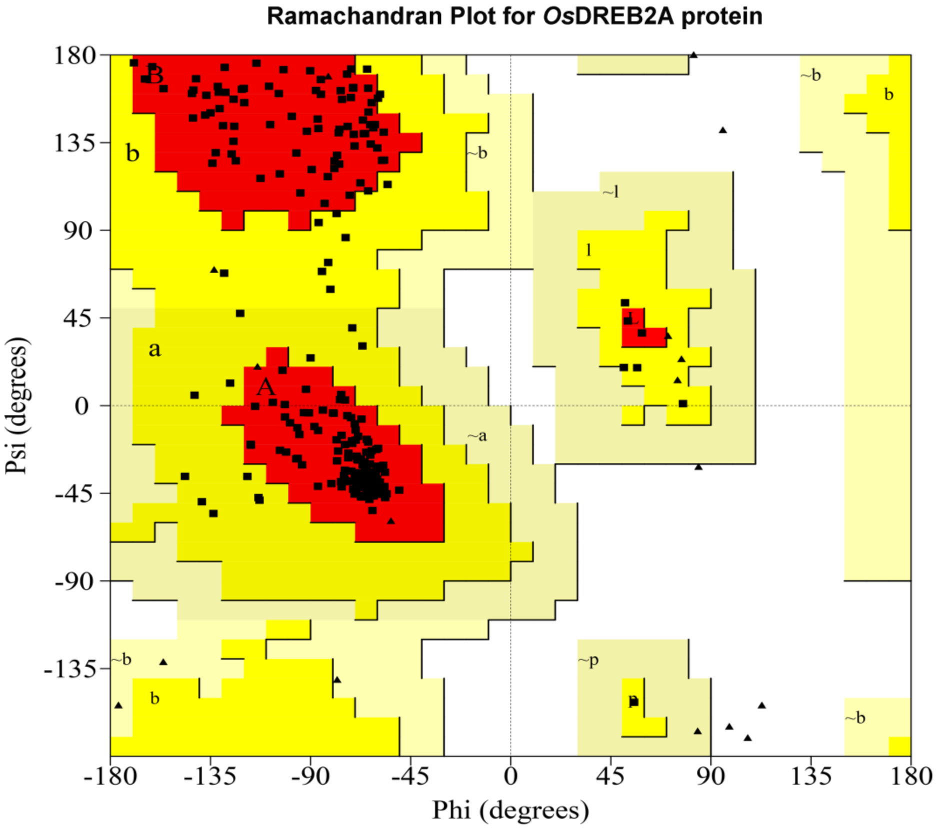


Plot statistics

| | | |
|---|---|---|
| Residues in most favoured regions [A,B,L] | 213 | 90.3% |
| Residues in additional allowed regions [a,b,l,p] | 23 | 9.7% |
| Residues in generously allowed regions [~a,~b,~l,~p] | 0 | 0.0% |
| Residues in disallowed regions | 0 | 0.0% |
|  | ---- | ------ |
| Number of non-glycine and non-proline residues | 236 | 100.0% |
| Number of end-residues (excl. Gly and Pro) | 2 | |
| Number of glycine residues (shown as triangles) | 22 | |
| Number of proline residues | 14 | |
|  | ---- |  |
| Total number of residues | 274 | |

**Supplementary Fig. S6: Ramachandran plot for *Os*DREB2A structure.**

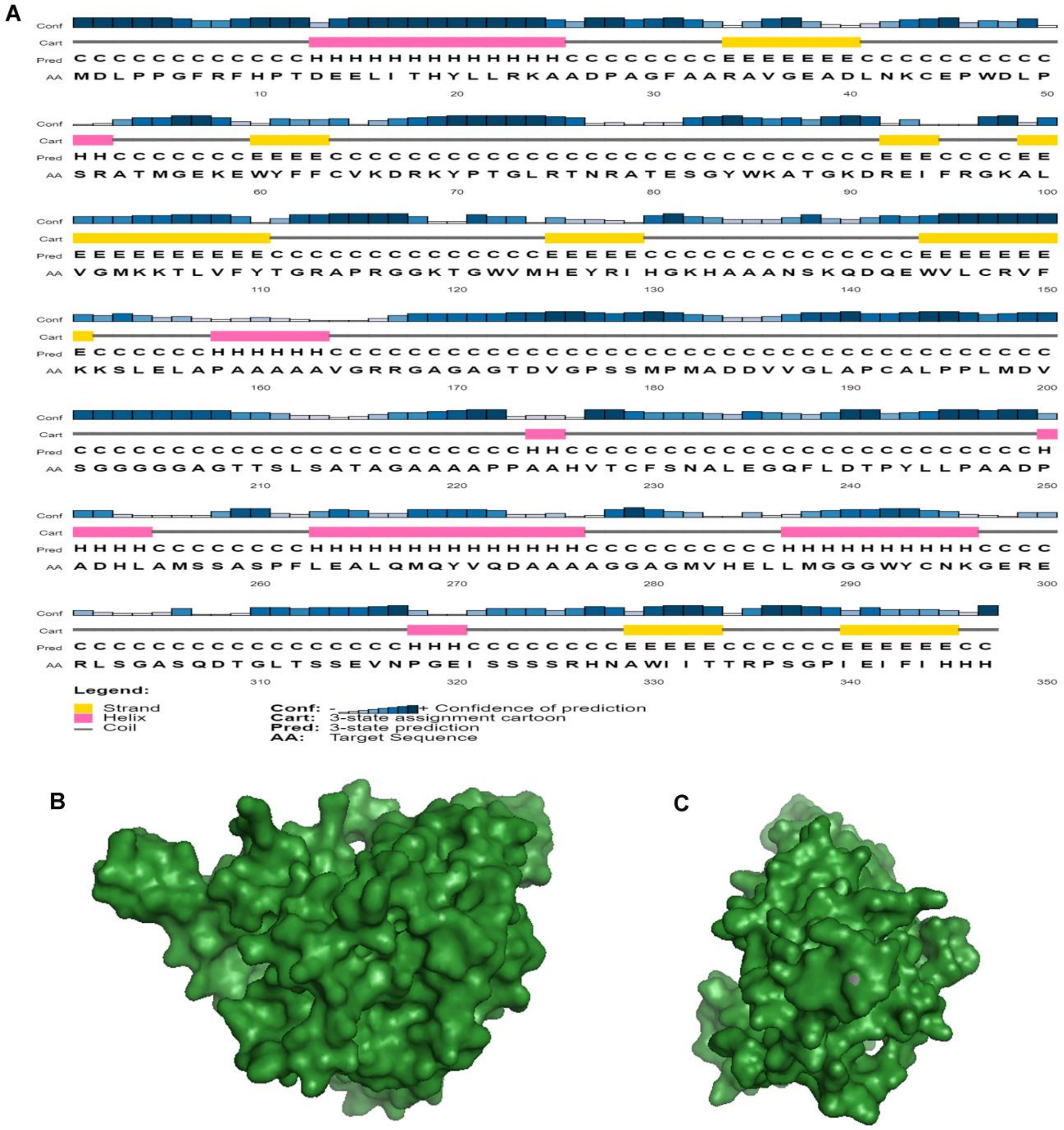


**Supplementary Fig. S7: Homology modelling of *Os*NAC1 protein.** (A) Fold prediction using PSIPRED, Surface view of the *Os*NAC1 protein in (B) side view and (C) top view.

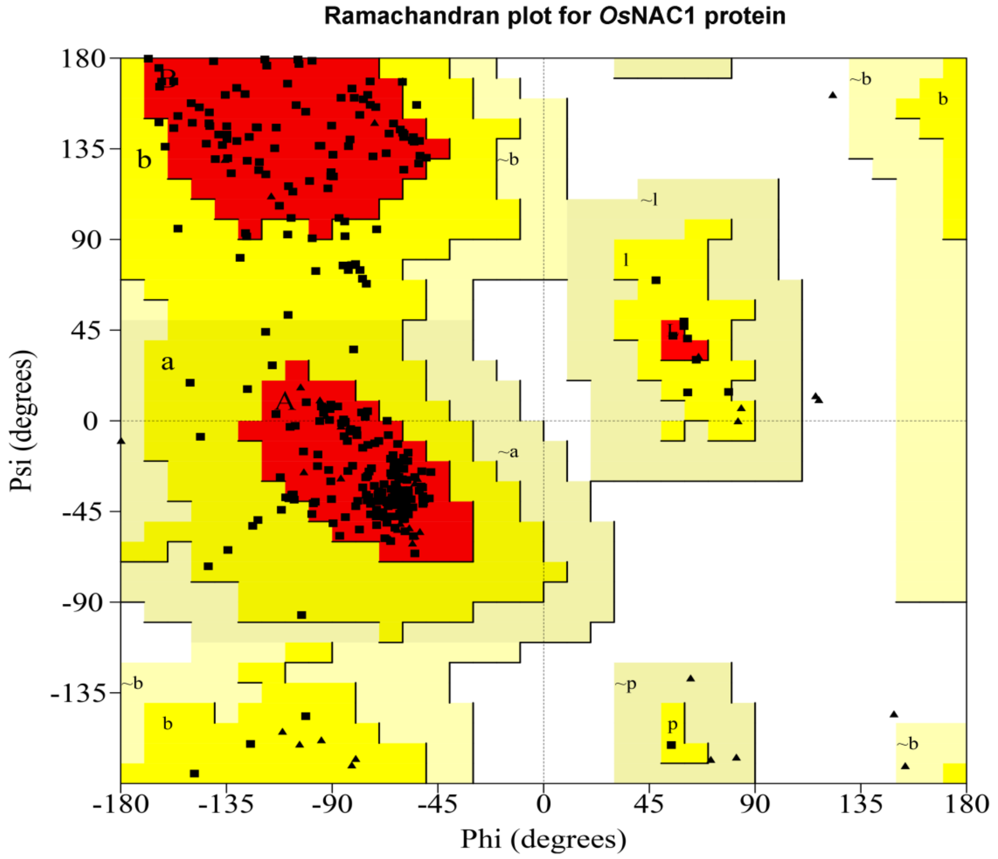


Plot statistics

| | | |
|---|---|---|
| Residues in most favoured regions [A,B,L] | 248 | 87.6% |
| Residues in additional allowed regions [a,b,l,p] | 35 | 12.4% |
| Residues in generously allowed regions [~a,~b,~l,~p] | 0 | 0.0% |
| Residues in disallowed regions | 0 | 0.0% |
|  | ---- | ------ |
| Number of non-glycine and non-proline residues | 283 | 100.0% |
| Number of end-residues (excl. Gly and Pro) | 2 | |
| Number of glycine residues (shown as triangles) | 39 | |
| Number of proline residues | 23 | |
|  | ---- |  |
| Total number of residues | 347 | |

**Supplementary Fig. S8: Ramachandran plot for *Os*NAC1 structure.**

**Sequence analysis for *Os*NAC1 protein**

```
MDLPPGFRFHPTDEELITHYLLRKAADPAGFAARAVGEADLNKCEPWDLPSRATMGEKEW
YFFCVKDRKYPTGLRTNRATESGYWKATGKDREIFRGKALVGMKKTLVFYTGRAPRGGKT
GWVMHEYRIHGKHAAANSKQDQEWVLCRVFKKSLELAPAAAAAVGRRGAGAGTDVGPSSM
PMADDVVGLAPCALPPLMDVSGGGGGAGTTSLSATAGAAAAPPAAHVTCFSNALEGQFLD
TPYLLPAADPADHLAMSSASPFLEALQMQYVQDAAAAGGAGMVHELLMGGGWYCNKGERE
RLSGASQDTGLTSSEVNPGEISSSSRHNAWIITTRPSGPIEIFIHHH
```

DNA-binding domain

*Os*SalT-binding domain

**Supplementary Fig. S9: Sequence analysis for *Os*NAC1 protein showing the DNA-binding and OsSalT-interacting sites.**

**A** **Sequence analysis for *Os*DREB2A protein**

MERGEGRRGDCSVQVRKKRTRRKSDGPDSIAETIKWWKEQNQKLQEENSSRKAPAKGSKK
GCMAGKGGPENSNCAYRGVRQRTWGKWVAEIREPNRGRRLWLGSFPTALEAAHAYDEAAR
AMYGPTARVNFADNSTDANSGCTSAPSLMMSNGPATIPSDEKDELESPPFIVANGPAVLY
QPDKKDVLERVVPEVQDVKTEGSNGLKRVCQERKNMEVCESEGIVLHKEVNISYDYFNVH
EVVEMIIVELSADQKTEVHEEYQEGDDGFSLFSY

**B** **Ubiquitination site prediction**

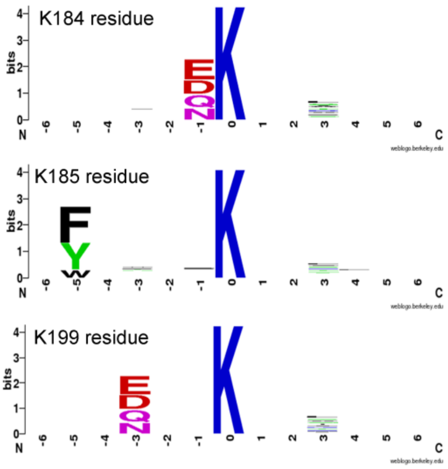


DNA-binding domain
Acidic residues in putative NRD domain
Serine/threonine residues in putative NRD domain
Ubiquitination site
AA *Os*SalT-binding domain
White font: SUMOylation site

**C** **SUMOylation site prediction**

| Position | Peptide | Score | Cutoff | P-Value | Type |
|---|---|---|---|---|---|
| **199** | VPEVQDVKTEGSNGL | 15.784 | 3.24 | 0.005 | SUMOylation Concensus |

**Supplementary Fig. S10: Sequence analysis for *Os*DREB2A protein.** (A) Protein sequence showing the DNA-binding, OsSalT-interacting, ubiquitination and SUMOylation consensus sites. (B) Ubiquitination site prediction. (C) SUMOylation site prediction.
