## Supplementary material for "Jacalin domain-containing protein *Os*SalT interacts with *Os*DREB2A and *Os*NAC1 to impart drought stress tolerance *in planta*": ESM_2

*****Authors for correspondence:

**Table S1: List of primers used.**

| Genbank Accession No | Gene name | Primers |
|---|---|---|
| XM_015774830 | *Osactin1* | Forward-5’CCGTCCTCCTGCTTGTTTCT3’ |
| | | Reverse: 5’TGGTACCCTCATCAGGCATC3’ |
| GQ339768.1 | *Ntactin* | Forward: 5’CCATCTACGAGGGTCATGCT3’ |
| | | Reverse: 5’ATCATGGATGGCTGGAAGAG3’ |
| U60097.2 | *Osdehydrin* | Forward: 5’ATGAGGGAGGAGCACAAGAC3’ |
| | | Reverse: 5’TCCATGATGCCCTTCTTCTC3’ |
| JN848806 | *OsSalT(partial)* | Forward: 5’GCCACCCAAGAAGCTGTTAG3’ |
| | | Reverse: 5’GTCTTGCAGTGGAATGCTGA3’ |
| AB049337 | *NtDehydrin* | Forward: 5’GAGGAAAAGCCAACTCATGC3’ |
| | | Reverse: 5’ACTGGTACAGCCGTGTCCTC3’ |
| GQ354807.1 | *NtLEA* | Forward: 5’AACCTGAGGCAAGCATCACT3’ |
| | | Reverse: 5’GTTGCCAATGACTGGAAGGT3’ |
| XM_016579794 | *NtSPS* | Forward: 5’GACAATGGAAGCCTGGGTTA3’ |
| | | Reverse: 5’TCTCCCCCATCTCAGTCATC3’ |
| NM_001325604 | *NtPI3K* | Forward: 5’AGTGAAGAGGTCCGTGCCTA3’ |
| | | Reverse: 5’CCCCAACTTCAACATGCTTT3’ |
| P16426 | *Bar* | Forward: 5’AAGCACGGTCAACTTCCGTA3’ |
| | | Reverse: 5’GAAGTCCAGCTGCCAGAAAC3’ |
| JN848806 | *OsSalT* | Forward: 5’GAATTCATGACGCTGGTGAAGATTGG3’ |
| | | Reverse: 5’GGATCCTCAAGGGTGGACGTAGATGC3’ |
| XM_026022985 | *OsDREB2A* | Forward: 5’ACTAGTTTCCCGCTCGATGGAGCG3’ |
| | | Reverse: 5’GGATCCCTAATAGGAGAAAAGGCTAAAC3’ |
| AB028180 | *OsNAC1* | Forward: 5’ACTAGTAGATGGACCTGCCGCCGG3’ |
| | | Reverse: 5’GGATCCCAACACAATCAATAATCGTGCC3’ |
| XM_016642058 | *NtRD22* | Forward: 5’GCTGTAGTTTGCCACAAGCA3’<br>Reverse: 5’ GAAGGAAATGGCAAACAGGA3’ |
| XM_016647041 | *NtADH1* | Forward: 5’GGAGGTGTTGACCGAAGTGT3’ |
| | | Reverse: 5’ CATGTACTTGCCCACCACAG3’ |
